# Reliability of neural entrainment in the human auditory system

**DOI:** 10.1101/2020.12.18.423536

**Authors:** Yuranny Cabral-Calderin, Molly J. Henry

## Abstract

Auditory stimuli are often rhythmic in nature. Brain activity synchronizes with auditory rhythms via neural entrainment, and entrainment seems to be beneficial for auditory perception. However, it is not clear to what extent neural entrainment in the auditory system is reliable over time – a necessary prerequisite for targeted intervention. The current study aimed to establish the reliability of neural entrainment over time and to predict individual differences in auditory perception from associated neural activity. Across two different sessions, human listeners detected silent gaps presented at different phase locations of a 2-Hz frequency modulated (FM) noise while EEG activity was recorded. As expected, neural activity was entrained by the 2-Hz FM noise. Moreover, gap detection was sinusoidally modulated by the phase of the 2-Hz FM into which the gap fell. Critically, both the strength of neural entrainment as well as the modulation of performance by the stimulus rhythm were highly reliable over sessions. Moreover, gap detection was predictable from pre-gap neural 2-Hz phase and alpha amplitude. Our results demonstrate that neural entrainment in the auditory system and the resulting behavioral modulation are reliable over time, and that both entrained delta and non-entrained alpha oscillatory activity contribute to near-threshold stimulus perception. The latter suggests that improving auditory perception might require simultaneously targeting entrained brain rhythms as well as the alpha rhythm.

## Introduction

Auditory stimuli, such as music and speech, are often (quasi-)rhythmic in nature. As such, one neural mechanism that has received much recent attention for its contribution to auditory perception is synchronization of brain activity to sound rhythms: *neural entrainment* (Obleser and Kayser, 2019). Neural entrainment is the process by which neural activity phase locks to sensory rhythms, and has been proposed to be a key mechanism for controlling sensory gain (Lakatos et al., 2008; Obleser and Kayser, 2019), attention, and parsing (Giraud and Poeppel, 2012) sensory information that is extended in time.

Neural entrainment has been described for different sensory modalities (Henry and Obleser, 2012; Mathewson et al., 2012; Ten Oever et al., 2017) and different species (Lakatos et al., 2016; Garcia-Rosales et al., 2018). In humans, low-frequency delta/theta activity synchronizes to the rhythms of speech and music, and synchronization success seems to be critical for successful auditory perception (Doelling et al., 2014; Doelling and Poeppel, 2015), in particular in noisy listening situations (Horton et al., 2013; Zion Golumbic et al., 2013; Brodbeck et al., 2020).

If neural entrainment does play a critical mechanistic role for auditory perception, *improving* synchrony between brain activity and stimulus rhythms should result in benefits for perception. In fact, recent work using transcranial electrical stimulation (TES) with alternating current (tACS) to interfere with entrainment reported significant modulation of speech comprehension and stream segregation (Riecke et al., 2015; Riecke et al., 2018; Wilsch et al., 2018; Zoefel et al., 2018; Cabral-Calderin and Wilke, 2020). However, there is uncertainty in tACS results (Erkens et al., 2020), and its efficacy for modulating neural entrainment is still under debate. This is at least in part because TES effects are state-dependent and variable across participants (Cabral-Calderin et al., 2016; Kasten et al., 2019), meaning that the same TES parameters might not be optimal for all participants, or even for one participant under all circumstances. In particular, successfully interfering with neural entrainment likely depends on targeting the right frequency (Stecher and Herrmann, 2018) and stimulus–brain phase relationship (van Bree et al., 2021), which may vary from person to person or from day to day. Here, we aimed to establish the reliability of neural entrainment over time. In doing so, our goal was to demonstrate the viability of an approach whereby individualized TES protocols can be based on entrainment signatures measured in a separate EEG session without the need for concurrent electrophysiological measures while the electrical stimulation is applied, which is very challenging.

Not all stimuli are rhythmic. Thus, multiple neural processing modes have been proposed: “rhythmic-mode” and “continuous-mode” processing (Schroeder and Lakatos, 2009; Henry and Herrmann, 2012), only the former of which relies on neural entrainment. Even in the case of rhythmic stimulation, entrainment lapses are associated with periods of high-amplitude alpha oscillations (Lakatos et al., 2016), and higher alpha amplitude has also been related to reduced entrainment-driven behavioral modulation (Henry et al., 2017). Taken together, these findings suggest that alpha activity and entrainment represent interacting neural strategies that can comodulate behavior. Here, in addition to examining the influence of entrained neural activity on auditory perception (Henry et al., 2014; Bauer et al., 2018), we took the influence of alpha activity into account in order to provide a more complete picture of the neural mechanisms underlying auditory perception in a vigilance task utilizing rhythmic stimuli.

Across two EEG sessions, participants detected silent gaps in ongoing frequency-modulated (FM) stimuli (**Fig. 1**, Henry and Obleser, 2012). We expected that the 2-Hz stimulus rhythm would entrain delta oscillations and, as such, gap detection would be modulated by the phase of both the stimulus and the entrained neural oscillation in which gaps occurred. We show that, while neural entrainment and stimulus-driven behavioral modulation are highly reliable across sessions, auditory perception cannot be exclusively explained by entrained low-frequency neural activity: both entrained delta phase and non-entrained alpha activity contribute to moment-by-moment variations in auditory perception.

**Figure 1.**
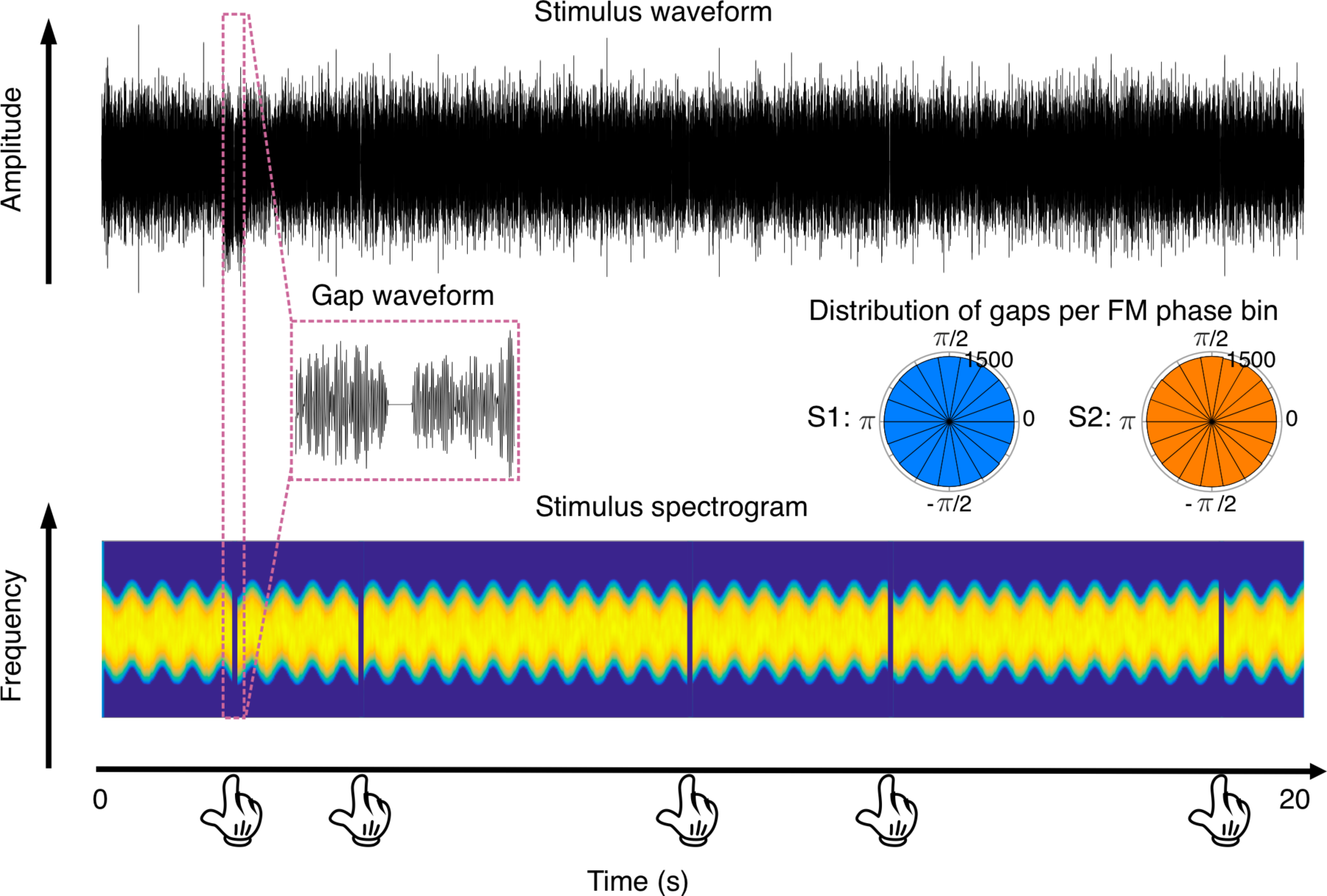
Auditory stimuli. Stimuli were 20-s long frequency modulated (FM) sounds whose frequency fluctuated rhythmically at 2 Hz (bottom), without periodic fluctuations in amplitude (top). Participants detected short silent gaps (middle left: Gap waveform) embedded in the sound in one of 18 possible phase bins, uniformly distributed around the 2 Hz FM cycle. Each sound had 3, 4, or 5 gaps. Participants responded with a button press each time they detected a silent gap. Circular histograms in the figure (middle right: Distribution of gaps per FM phase bin) show the distribution of gaps per phase bin across participants, separated by session. S1: session 1; S2: session 2. Hand icon was downloaded from https://www.stockio.com/.

## Materials and Methods

### Main experiment

#### Participants

Forty-one healthy participants took part in the study. Three participants were excluded from further analysis due to noisy EEG data (1 participant) and poor task performance (i.e., detection rate < 0.25, 2 participants). Results presented in this manuscript include data from 38 participants (21 females, four left-handed, mean age = 26.03 with SD = 4.6 years old). Each participant took part in two sessions separated by 2–42 days (median: 7 days). All participants self-reported normal-hearing and normal or corrected-to-normal vision. All participants were either native German speakers (n = 37) or spoke German with high proficiency (n = 1). At the time of the experiment no participant was taking medication for any neurological or psychiatric disorder.

Participants received financial compensation for their participation in the study. Written informed consent was obtained from all participants. The procedure was approved by the Ethics Council of the Max Planck Society and in accordance with the declaration of Helsinki.

#### Stimuli

Auditory stimuli were generated by MATLAB software at a sampling rate of 44,100 Hz. Stimuli were 20-s long complex tones frequency modulated at a rate of 2 Hz and with a center-to-peak depth of 67% (**Fig. 1**). The center frequency for the complex carrier signals was randomly chosen for each stimulus within the range of 1000-1400Hz. The complex carrier comprised 30 components sampled from a uniform distribution with a 500-Hz range. The amplitude of each component was scaled linearly based on its inverse distance from the center frequency; that is, the center frequency itself was the highest-amplitude component, and component amplitudes decreased with increasing distance from the center frequency. The onset phase of the stimulus was randomized from trial to trial, taking on one of eight values (0, π/4, π/2, 3π/4, π, 5π/4, 3π/2, 7π/4) with the constraint that each trial would always start with a phase different from its predecessor. All stimuli were rms-amplitude normalized. Three, four, or five silent gaps were inserted into each 20-s stimulus (gap onset and offset were gated with 3-ms half-cosine ramps) without changing the duration of the stimulus. Each gap was chosen to be centered in 1 of 18 equally spaced phase bins into which each single cycle of the frequency modulation was divided. No gaps were presented either in the first or the last second of the stimulus. A minimum of 1.5 s separated consecutive gaps.

#### Procedure

The experiment was conducted in an electrically shielded and acoustically isolated chamber and under normal-illumination conditions. Sound-level thresholds were determined for each participant according to the method of limits. All stimuli were then presented at 55 dB above the individual hearing threshold (55 dB sensation level, SL).

Gap duration was individually adjusted to detection-threshold levels using an adaptive-tracking procedure comprising two interleaved staircases and a weighted up-down technique with custom weights. During this procedure, participants detected a gap within a 4-s FM sound. Except for the duration, the sound had the same characteristics as in the main experiment. The descending staircase started with a gap duration of 150 ms and the ascending staircase started with a gap duration of 1 ms. If the participant detected the gap, gap duration was decreased by some percent (5% for 10 ms ≤ gaps ≤ 35 ms, 20% for 35 ms < gaps ≤ 70 ms, or 50% for 70 ms < gaps || gaps < 10 ms) in the following trial of the current staircase. On the contrary, if the participant did not detect the gap, gap duration was increased by some percent (following the same convention as before) of the current gap duration, in the following trial of the current staircase. Each staircase ended when four reversals occurred in a span of six trials. The mean final gap duration across the two staircases was chosen as the gap size for that participant in the main task. This resulted in gap durations ranging from 11–28 ms across subjects and sessions (session 1: mean = 17.26 ms, SD = 3.42 ms, range = 12–28 ms; session 2: mean = 15.55, SD = 2.78, range = 11-21 ms).

Before starting the main experiment, participants performed practice trials to make sure they understood the task. For the main experiment, EEG was recorded while listeners detected gaps embedded in the 20-s long FM stimuli. Listeners were instructed to respond as quickly as possible when they detected a gap via button-press. Overall, each listener heard 216 stimuli (27 per starting phase). The number of gaps per stimulus was counterbalanced (72 stimuli each included 3 gaps, 4 gaps, and 5 gaps) for a total of 864 gaps. For each of the 18 FM-phase bins, 48 gaps were presented. The number of phase bins was chosen to balance having a good sampling resolution and a high number of trials per condition without making the task too long. This combination of phase bins and trials exceeds the recommendation for estimating the sinusoidal modulation of behavior with high sensitivity (Zoefel et al., 2019). Including the EEG preparation, each experimental session lasted about 3 hours.

### Data Acquisition and Analysis

#### Behavioral data

Behavioral data were recorded online by MATLAB 2017a (MathWorks) in combination with Psychtoolbox. Sounds were presented at a rate of 44.1kHz, via an external soundcard (RME Fireface UCX 36-channel, USB 2.0 & FireWire 400 audio interface) using ASIO drivers. Participants listened to the sounds via over-ear headphones (Beyerdynamic DT-770 Pro 80 Ohms, Closed-back Circumaural Dynamic Diffuse field equalization Impedance: 80 Ohm SPL: 96 dB Frequency range: 5–35,000 Hz). Button presses were collected using a Cedrus response pad (RB-740). Hits were defined as button-press responses that occurred no earlier than 100 ms and no later than 1.5 s after the occurrence of a gap (Henry et al., 2014). This window was chosen because the upper limit represents the minimum separation between gaps and therefore the response can easily be assigned to a specific gap. Decreasing this window to maximum 1 s after gap onset produced qualitatively similar results. Hit rates and RTs were calculated separately for each of the 18 FM-phase bins. To estimate the FM-induced sinusoidal modulation of gap detection behavior, a cosine function was fitted to hit rates as a function of FM phase for each participant and session. From the fitted function, the amplitude parameter quantifies the strength of behavioral modulation by 2-Hz FM phase, while the phase parameter indexes the FM-stimulus–brain lag. Significance of the sinusoidal modulation was tested using a permutation approach, whereby 1000 surrogate datasets were created for each participant and session by shuffling the single-gap accuracy values (0,1) with respect to their stimulus-condition labels. Cosine functions were also fitted to the surrogate datasets. Gap detection was considered to be sinusoidally modulated for each participant if the individual amplitude value was higher than the 99.9 percentile of the distribution of amplitude values from the surrogate data, corresponding to p < 0.05, *Bonferroni* corrected for 76 comparisons -i.e., 38 subjects x 2 sessions). Optimal FM phase was defined as the FM-stimulus phase with highest hit rate in the fitted function.

We aimed to test the influence of FM phase on gap detection while simultaneously testing and controlling for stimulus characteristics, as well as the passage of time. To do this, we fit mixed-effects logistic regression models using the MATLAB function ‘*fitglme*’. Each model was fitted using a binomial distribution, logit link function, and Laplace fitting method. Different models were tested evaluating whether gap detection could be predicted as a function of 1) FM phase, 2) stimulus center carrier frequency (1000–1400Hz), 3) stimulus carrier at gap onset, 4) global time (1–864, indicating the gap position within the whole experiment), 5) local time (1–12 equally spaced time bins, excluding the first and last second of the 20-s stimulus, indicating gap location within the stimulus), 6) cochlear entropy (see below) and the interactions of 7) FM phase * stimulus carrier frequency, 8) FM phase * global time and 9) FM phase * local time. Each predictor, excepting interaction terms, was modeled as a fixed factor plus a random intercept and slope to estimate the group-level effect while allowing for inter-individual variability in the predictor effect size. Due to computational limitations, interaction terms were modelled only with fixed terms. All predictors were transformed to z-scores prior to model fitting. Ten different models were fitted, each including a different combination of regressors, and the best model was selected using the Akaike’s information criterium (AIC, **Table 2**) and the Likelihood Ratio Test (LRT). All predictors involving circular data (phase angles) were linearized by calculating their sine and cosine. The separate beta estimates for the sine (sinb) and cosine (cosb) of the phase predictor were recombined into a single beta corresponding to the amplitude of the phasic behavioral modulation:

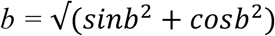

Cochlea-scaled spectral entropy is an index of spectral change and represents the relative unpredictability of acoustic signals (Henry et al., 2016; Teng et al., 2018). Since our perceptual systems are more driven to detect change than stability, it is possible that the instantaneous rate of frequency change at gap onset could affect gap detection. For calculating cochlea-scaled entropy (Stilp and Kluender, 2010; Henry et al., 2016; Teng et al., 2018), each 20-s auditory stimulus was divided into 16 ms “slices”. Each slice was filtered by a bank of 15 symmetrical rounded-exponential filters with center frequencies equally spaced between 300 and 2400 Hz in equivalent-rectangular-bandwidth (ERB) units. The result was a 15-bin frequency histogram where the height of each bar was the amplitude of the corresponding filter output. Pairwise Euclidean distances were then calculated for frequency histograms belonging to neighboring slices. For each gap, cochlea-scaled entropy was finally computed by calculating the Euclidean distance values averaged in a 27-ms time window right before gap onset. This duration was chosen because it corresponds to the duration of each phase bin.

To test the classification significance of the winning model, 1000 surrogate datasets were created for each session by shuffling the single-gap accuracy values (0,1) for each participant while keeping all condition labels the same. The same mixed-effects models were fitted to the surrogate datasets and the response operator characteristic (ROC) curve and the area under the curve (AUC) for the winning model and each surrogate model were computed. The winning model was considered to be significant if its AUC value fell above the 95 percentile of the distribution of AUC from the surrogate models (p < 0.05).

### Electroencephalogram data

The EEG was recorded with an actiCAP active electrode system in combination with Brainamp DC amplifiers (Brain Products GmbH). The electrode system included 64 Ag–AgCl electrodes mounted on a standard cap, actiCAP 64Ch Standard-2 (Brain Products GmbH). Signals were recorded continuously with a passband of 0.1 to 1000 Hz and digitized at a sampling rate of 1000 Hz. For recording, the reference electrode was placed over FCz and the ground electrode over AFz. For better stimulus marking, in addition to standard EEG triggers from the LPT port, stimulus markers were also sent via soundcard and collected in the EEG using a Stimtrak (Brain Products GmbH). Electrode resistance was kept under 20 kΩ. All EEG data were analyzed offline by using Fieldtrip software (www.ru.nl/fcdonders/fieldtrip; version 20200130) and custom MATLAB scripts.

Two different preprocessing pipelines were implemented. One was tailored to assess entrainment characteristics and reliability, and focused on the complete 20-s stimulus periods; the second pipeline was tailored to test the effect of pre-target (pre-gap) activity on gap detection, and focused on the periods around each gap’s occurrence. In the first preprocessing pipeline, the continuous EEG data were high-pass filtered at 0.6 Hz. Filtered data were then epoched into 21.5-s trials (1 s before stimulus onset and 0.5 s after stimulus offset). The trial data were low-pass filtered at 80 Hz and the 50-Hz line noise was removed using discrete Fourier transform (dft) with spectrum interpolation as implemented in Fieldtrip. Data were re-referenced to the average reference. Extreme artifacts were removed based on visual inspection. Noisy electrodes were then interpolated (1 electrode in 3 participants and 2 electrodes in one participant). Eye-blinks, muscle, heartbeat, and remaining line noise or faulty contact artifacts were removed using ICA. Next, data were low-pass filtered to 30 Hz and trials for which the range exceeded 200 µV were automatically removed. If more than 30% of the trials had to be removed because of artifacts, the participant was removed for further analysis (1 participant). Preprocessed data were resampled to 500 Hz.

The second preprocessing pipeline included the same steps excepting the initial high-pass filter. To maximize comparability with the first pipeline, all the same trials and ICA components that were identified based on the first pipeline were removed in the second pipeline. After all preprocessing steps and before resampling, 3-s long trials were defined around each gap onset (i.e., 1.5 s before and 1.5 s after gap onset). Trials exceeding a range of 200 µV were excluded and data were resampled to 500 Hz.

#### Frequency and time-frequency analysis of full-stimulus periods

Full-stimulus epochs were analyzed in the frequency and time-frequency domains to examine brain responses entrained by the 2-Hz stimulation. Since the starting phase of the FM stimulus was randomized from trial to trial, before conducting frequency-domain analyses, single-trial brain responses were realigned so that the FM stimulus phases would be perfectly phase-locked across trials after the realignment. A fast Fourier transform (FFT) was performed on the trial-averaged time-domain data at each electrode, after multiplication with a Hann window. Evoked amplitude in each frequency band was calculated as the absolute value of the complex output, while the phase angle of the complex FFT output at 2 Hz provided an estimate of stimulus–brain phase lag. An FFT was also applied on each single trial, and the resulting single-trial amplitude spectra were averaged over trials as an indicator of total amplitude of neural activity that was not necessarily phase-locked to the stimulus. Inter-trial phase coherence (ITPC) was calculated as the resultant vector length of phase angles from the complex FFT output across trials separately for each frequency and electrode. In addition, the single-trial time-domain data were submitted to a time–frequency analysis by using the Fieldtrip-implemented version of the Wavelet approach using Fourier output. Here, wavelet size varied with frequency linearly from three to seven cycles over the range from 1 to 15 Hz. The resulting complex values were used to estimate time-resolved ITPC for each channel separately.

Nonparametric Wilcoxon signed-rank tests were conducted to statistically test spectral amplitudes and ITPC at frequencies of interest (2 Hz, 4 Hz, alpha). For each condition, participant, and session, data were averaged over all channels (and over time for time-resolved ITPC) and amplitudes/ITPC of the two target frequencies (2 Hz and 4 Hz) were then tested against the average amplitude/ITPC of the neighboring ±8 frequency bins (0.16Hz) (Henry and Obleser, 2012; Bauer et al., 2018). In the case of alpha amplitude, data were averaged across all bins including frequencies between 7-12 Hz and were tested against the average amplitude of the neighboring ±100 frequency bins (2 Hz).

Based on the topography of the 2/4 Hz and alpha (7-12 Hz) amplitude spectra, further analyses involving the extraction of ITPC and/or amplitude values were done in two separate clusters of electrodes including *F3, Fz, F4, FC1, FC2, C3, Cz, C4, F1, F2, FC3, FC4, C1, C2* for 2/4 Hz activity and *P8, P6, P4, P2, Pz, P1, P3, P5, P7, PO9, PO10, PO8, PO4, POz, PO3, PO7, O1, Oz, O2* for alpha activity. First, ITPC or amplitude values were extracted from each electrode and then averaged over electrodes within the relevant cluster.

### Pre-gap activity

Before analysis of pre-gap activity, single-trial time-domain data around the gap period (1.5 s before and 1.5 s after) were detrended (using linear regression). It is possible that the smearing of the evoked response back into pre-stimulus period by wavelet convolution could produce spurious pre-stimulus phase effects. To minimize this, gap-evoked responses were removed from the post-stimulus period by multiplication with half of a Hann window that ranged between 0 and 50 ms after gap onset and was zero thereafter (Henry et al., 2014). To estimate neural amplitude and phase, the time-domain data were submitted to a wavelet convolution yielding complex output as implemented in Fieldtrip. Wavelet size varied with frequency linearly from three to seven cycles over the range from 1 to 15 Hz with 10-ms temporal resolution. Alpha amplitude as well as 2-Hz amplitude were averaged within the 100-ms time window preceding gap onset. Neural phase was computed as the phase angle of the complex output for the specific frequency. Time windows for extracting pre-gap 2-Hz and alpha phases were adjusted to include 1/5 of a cycle of the relevant frequency, i.e., 100 ms preceding gap onset for 2-Hz phase and 22 ms for alpha phase (assuming center frequency 9 Hz). Since different electrodes can have different phase profiles, computing circular means over different electrodes can lead to phase cancellation and an imprecise estimation of phase effects on gap perception. To circumvent this problem, phase values were realigned to a group reference per electrode, prior to calculating the circular mean across electrodes within the relevant electrode cluster (fronto-central cluster for 2-Hz phase estimation and parieto-occipital cluster for alpha phase estimation). This way we hoped to minimize inter-electrode variability within the cluster, while maintaining inter-participant and inter-session variability. First, trials were binned according to their pre-gap 2-Hz/alpha phase and hit rates calculated for each bin. Cosine functions were fitted to binned hit-rate data for each single participant and session and the optimal phase for gap detection estimated. The group phase references (∅_*ref*_) were estimated for each electrode by computing the circular mean optimal phase over sessions and participants. For each electrode, single trial phase (∅_(*tr*)_) values were then realigned (∅*r*_(*tr*)_) as:

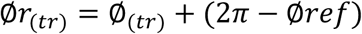

The influence of pre-gap 2-Hz phase and amplitude, alpha phase and amplitude, and their interactions on gap detection was evaluated using mixed-effects logistic regression models, similar to the behavioral analysis, using the MATLAB function *‘fitglme’*, specifying the distribution as binomial, the link function as logit, and the fitting method as Laplace. Fourteen different logistic regression models were fitted to the data including different combinations of the regressors, and the best model was chosen based on the AIC value (**Table 1**) and the LRT. Phase regressors were linearized by computing their sine and cosine. The phase–phase interaction between 2-Hz phase and alpha phase was assessed using two predictors, i.e., the product of the sines and the product of the cosines. The interaction between 2-Hz phase and alpha amplitude was assessed also using two predictors, i.e., the product of alpha amplitude and the sine and the cosine of 2-Hz phase. All predictors were transformed to z-scores prior to model fitting. To test the significant classification performance of the winning model, 1000 surrogate datasets were created for each participant and session by shuffling the single-gap accuracy values (0,1) across trials while preserving trial correspondence across regressors. The same regression models were fitted to the surrogate datasets. ROC and AUC values were computed for both the winning model and the surrogate models. The winning model was considered to be significant if its AUC value fell above the 95 percentile of the distribution of AUC from the surrogate models (p < 0.05).

To investigate optimal 2-Hz phase for gap detection, trials were sorted into 18 equally spaced phase bins based on realigned pre-gap 2-Hz phase angle, and hit rates were calculated for each bin. Optimal 2-Hz, defined as the phase with highest detection rate, was estimated based on separate cosine fits to hit rates binned by either delta phase.

#### Estimating 2-Hz phase–alpha amplitude coupling

To investigate 2-Hz phase–alpha amplitude coupling, generalized linear mixed-effects models were used to predict pre-gap alpha amplitude from 2-Hz pre-gap phase (its sine and cosine). The model included fixed terms as well as random intercept and slope. Individual beta estimates for the effect of 2-Hz phase on alpha amplitude (here interpreted as cross-frequency coupling) were recombined into a single beta for the phase effect, as described above. To test for statistical significance of the coupling, 1000 surrogate datasets were created for each participant and session by shuffling the trial order for the response variable (alpha amplitude) while keeping the order fixed for the predictor (2-Hz phase). The same generalized linear mixed effects models were fitted to each surrogate dataset and the beta coefficients were estimated just as described for the real data. Cross-frequency coupling was considered to be significant if the beta coefficients from the real model fell above the 95 percentile of the distribution of beta coefficients from the surrogate models (p < 0.05).

### Questionnaires

To evaluate musical skills, all participants from the main experiment completed the Goldsmiths Musical Sophistication Index (Gold-MSI) (Mullensiefen et al., 2014). Scores were computed using the documents and templates provided in https://www.gold.ac.uk/music-mind-brain/gold-msi/download/. Accordingly, individual scores were used to calculate the general sophistication index as well as scores on five main factors: active engagement, perceptual abilities, musical training, emotions, singing abilities.

### Statistical Analysis

Unless otherwise specified, correlation analyses between linear variables were done using Spearman rank correlation coefficient. In case of circular data, circular–circular or circular–linear correlations were computed using the respective functions in the circular statistics toolbox for MATLAB (Berens, 2009). Significant differences between sessions were evaluated using Wilcoxon signed-rank tests. Significant p-values were corrected using *Bonferroni* method. All statistics values in the text are rounded to max 3 decimal points.

### Control study

Twenty-five participants took part in the control experiment (seventeen females, mean age 26.08 (SD = 4.62)). Seventeen of them received financial compensation and signed written informed consent (10 also participated in the main study). The other eight participants were colleagues in our research group/institute (including one author-YCC), and participated voluntarily without compensation.

Unless otherwise specified, stimuli and procedures were the same as for the main experiment. In contrast to the main experiment, gap thresholds were not individually estimated, but gap duration was fixed at 16 ms (mean threshold in session 2 in the main experiment) for all participants. FM stimuli were created as for the main experiment but modulated at four different frequencies: 1.5, 2, 2.5, and 3 Hz. Gaps could be presented at 15 different bins of the FM cycle. Subjects heard 224 stimuli (56 per FM) for a total of 932 gaps (233 per FM rate). Stimuli were presented in 8 different blocks (2 per FM rate) of 28 stimuli each. Each block comprised only one FM rate but the FM order was randomized within session and across participants. The number of gaps per stimulus (3, 4, or 5) was randomized. For each of the 15 FM-phase bins*FM rate combination, 14-16 gaps were presented.

Hits were defined as button-press responses that occurred no earlier than 100 ms and no later than 1.5 s after the occurrence of a gap. Hit rates were calculated separately for each of the 15 FM-phase bins and each FM rate. To estimate the FM-induced sinusoidal modulation of gap detection behavior, a cosine function was fitted to hit rates as a function of FM-phase for each participant. For each participant and FM rate, the mean hit rate (i.e., fitted intercept), the fitted amplitude parameter, and the optimal phase (same definition as in the main experiment) were estimated. The effect of FM rate on hit rates and amplitude parameters was tested using separate one-way Analyses of Variance (ANOVAs). To test whether the concentration of optimal stimulus phases changed with FM rate, resultant vector lengths were computed for distributions of individual optimal phases, weighted by individual amplitude parameters, separately for each FM rate. A linear model was fitted to the data, predicting resultant vector length from FM rate and an intercept. The model was fitted to the data using the Matlab function *“fitglm”* and statistical significance was estimated using permutation tests. For comparison, the test distribution was created by computing 1000 resultant vector lengths calculated with the individual optimal phases while shuffling the FM rate information at the individual level. Data and Matlab codes will be made publicly available in https://edmond.mpdl.mpg.de/imeji/collection/9UoanSkXciMEK_v?q=afterpublication.

## Results

In two different sessions, EEG activity was recorded while listeners detected silent gaps embedded in 20-s long complex tones that were frequency modulated at 2 Hz (**Fig. 1**). Based on previous literature, we predicted that delta oscillations would be entrained by the 2-Hz FM. We aimed to quantify how reliable entrainment would be across sessions, both in terms of entrainment strength and in terms of the phase relationship between stimulus and brain.

### Auditory entrainment to FM sounds has high inter-session reliability

We evaluated entrainment using four converging analyses: (1) total and (2) evoked amplitudes of EEG data, (3) inter-trial phase coherence (ITPC) across the full epoch based on complex Fourier output, and (4) in a time-resolved way based on the output of a wavelet convolution (see Methods).

For evoked spectra, we observed high-amplitude peaks at the stimulus FM frequency and its first harmonic (2 Hz and 4 Hz respectively, **Fig. 2a**), consistent with neural tracking of the rhythm of the FM stimulus (Henry and Obleser, 2012; Henry et al., 2014). Relatively high spectral amplitude was also observed in the alpha frequency band (7-12 Hz). For all further analyses on alpha activity, we considered activity between 7 Hz and 12 Hz, because this frequency range best captured the observed range of alpha evoked amplitude (**Fig. 2a**). Evoked amplitudes for 2 Hz, 4 Hz, and alpha frequencies were significantly different than the average amplitude of the neighboring frequency bins (all z ≥ 5.344, p ≤ 5.452e-07, see Methods) in both sessions. Total amplitude spectra showed high amplitude in alpha frequency band, while 2-Hz and 4-Hz amplitudes were less visible in the spectra due to high 1/f power (**Fig. 2b**). Nevertheless, compared to the average amplitude of the neighboring frequency bins, 2 Hz, 4 Hz, and alpha amplitudes were also significant in the total amplitude spectra (all z ≥ 4.445, p ≤ 5.275e-05). Moreover, the ITPC analysis showed clear peaks at 2 Hz and 4 Hz, again suggesting neural tracking at the stimulus FM frequency and its first harmonic (**Fig. 2c**). In both sessions, ITPC at 2 Hz and 4 Hz was significantly different than the neighboring frequency bins (same neighboring frequency bins as defined for the evoked amplitude analysis, all z ≥ 5.330, p ≤ 3.938e-07). Finally, time-resolved ITPC at 2 Hz and 4 Hz, averaged over electrodes and time was significantly different than for the neighboring frequency bins (±8 frequency bins/0.81Hz, all z ≥ 5.069, p ≤ 1.604e-06, **Fig. 2c**, right). FM-stimulus-evoked amplitude at 2 Hz (and also at 4Hz, data not shown) was mostly observed in a fronto-central cluster including electrodes F3, Fz, F4, FC1, FC2, C3, Cz, C4, F1, F2, FC3, FC4, C1 and C2 (**Fig. 2a**, insets). Therefore, all further analyses involving 2 Hz and 4 Hz frequencies were done first independently by electrode and then averaged over this subset of electrodes; the one exception was the examination of neural phase effects of performance, where averaging over electrodes can lead to phase cancellation (see below). The same procedure was followed for all further analyses involving alpha frequencies but data were extracted from the cluster of electrodes exhibiting the highest stimulus-evoked alpha amplitude, i.e., P8, P6, P4, P2, Pz, P1, P3, P5, P7, PO9, PO10, PO8, PO4, POz, PO3, PO7, O1, Oz, O2 (**Fig. 2a**, insets).

**Figure 2.**
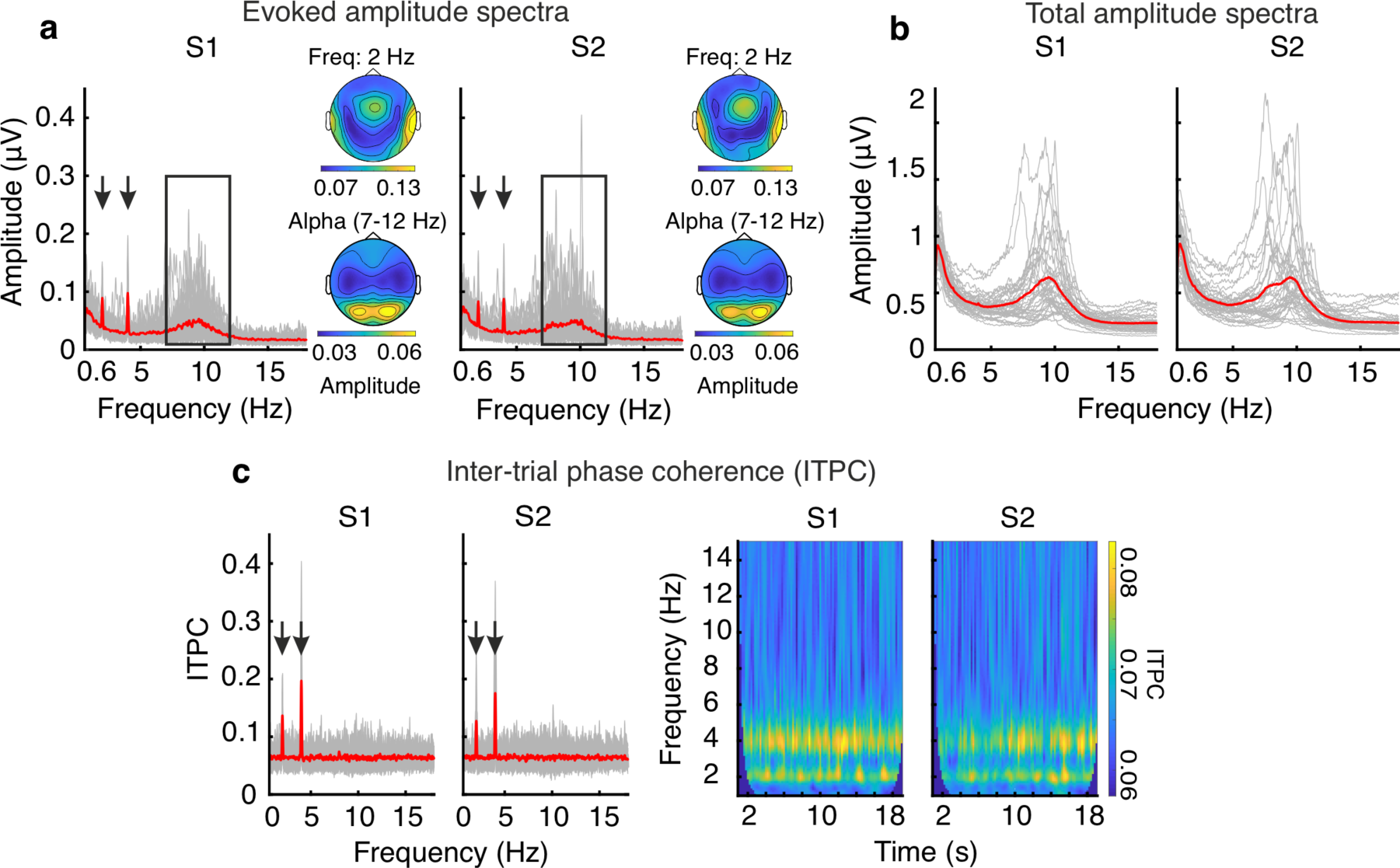
Neural entrainment to 2-Hz FM stimulus. (a) Evoked amplitude spectra from the fast Fourier transform (FFT) of the time-domain EEG signal. Red solid lines indicate the group average spectrum, gray lines show single participants’ spectra, averaged over all electrodes. Inset plots show the topography for 2 Hz and alpha amplitude spectra averaged across participants, separately for session 1 and session 2. (b) Total amplitude spectra. The spectra were computed independently for each trial and then averaged over trials for each subject. Red solid lines indicate the group average spectrum, gray lines show single participants’ spectra, averaged over all electrodes. (c) Inter-trial phase coherence (ITPC) indicating the degree of phase clustering across trials for each frequency (left), averaged over all electrodes. Gray lines show individual values and red lines show group average. (right) ITPC shown over time, again averaged across participants and all electrodes. S1: session 1; S2: session 2. Arrows in (a) and (b) indicate the peaks in amplitude and ITPC at the 2-Hz FM stimulus frequency and its first harmonic. Rectangles in (a) indicate the frequency range considered for further alpha analyses.

Moving a step past previous literature, we next asked whether FM-induced entrainment is reliable over time by correlating the amplitude of the stimulus-evoked activity at 2 Hz, 4 Hz, and in the alpha frequency band, as well as the stimulus-brain lag (i.e., the phase angle of the 2-Hz complex output of the FFT calculated for the full stimulus epoch) across sessions. Inter-session correlations were high and significant for evoked amplitudes at 2 Hz (rho = 0.701, p = 2.238e-06), 4 Hz (rho = 0.764, p = 2.846e-07), and in the alpha band (rho = 0.738, p = 5.962e-07; **Fig. 3a**). Significant difference between sessions was observed for 4-Hz amplitude (session 1 vs session 2: z = 2.632, p = 0.026, *Bonferroni* corrected) but not for the other frequencies (all |z| ≤ 1.429, p ≥ 0.153, uncorrected). Despite individual variability, stimulus–brain phase lags were not uniformly distributed as reported in previous work, but were significantly phase clustered for both sessions (Rayleigh test, session 1: z = 23.004, p = 9.513e-13; session 2: z = 16.087, p = 1.559e-08, **Fig. 3b**). Moreover, phase lags were reliable across sessions as indexed by the high circular–circular correlation (rho = 0.624, p = 0.003) and a circular distance between sessions clustered around zero (V-Test for nonuniformity with known mean direction, v = 30.981, p = 5.905e-13). Taken together, amplitude spectra and phase lags suggested that neural entrainment to FM stimuli is reliable across sessions, which is the first vital prerequisite for targeted interventions of auditory-cortex neural oscillations.

**Figure 3.**
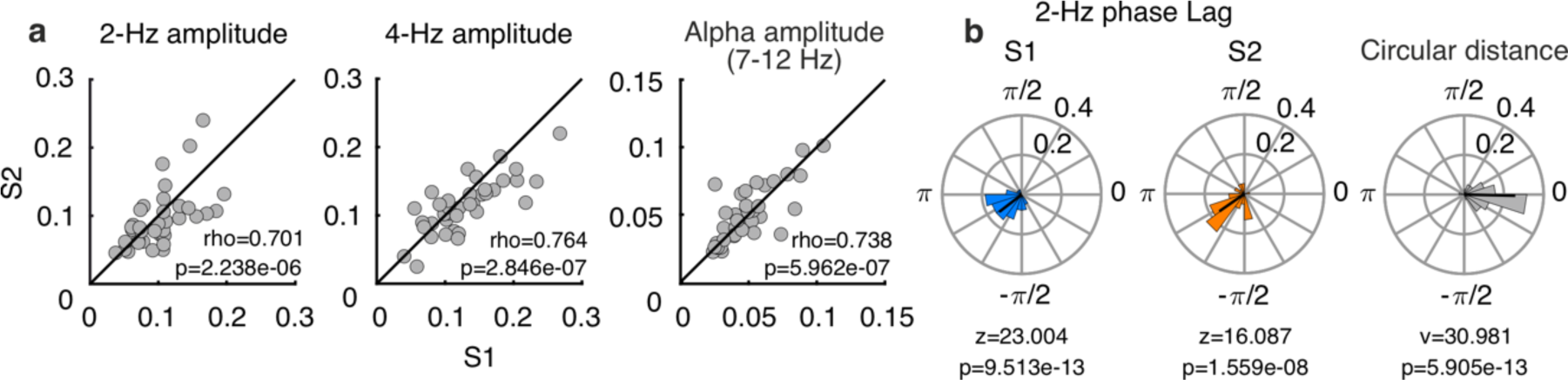
Reliability of neural entrainment. (a) Inter-session correlation of 2 Hz (left), 4 Hz (middle) and alpha (right) amplitudes. Correlation coefficients and associated p-values are given in each plot. Each dot represents a single participant. Solid black lines represent the diagonal for each graph. (b) Circular histograms show neural phase lag relative to the 2-Hz FM stimulus for session 1 (blue, left) and session 2 (orange, middle). Circular distance between phase lags in the different sessions is shown in the circular histogram in the right. Z-values and associated p-values from the Rayleigh test are given in each session plot and the v and p-value from the V-test is given for the circular-distance between sessions. S1: session 1; S2: session 2.

### Stimulus-induced behavioral modulation is sinusoidal and shows high inter-session reliability

While listening to the FM sounds, participants responded with a button press each time they detected a silent gap. Each 20-s long stimulus contained three, four, or five gaps. Gaps were distributed uniformly around the 2-Hz FM cycle in 18 possible phase positions (**Fig. 1**). A response was considered to be a “hit” if a button press occurred within a window of 0.1-1.5 s after gap onset.

We hypothesized that, as a consequence of the stimulus-induced entrainment of delta oscillations, hit rates for gap detection would be sinusoidally modulated by the FM stimulus phase. To go a step past previous studies, we also asked whether this modulation is reliable across sessions. While several studies have focused on analyzing oscillatory modulation of perception at the group level by aligning single-participant data to the phase with best or worst performance (see (Zoefel et al., 2019) for recommendations), here we took a different approach and investigated whether the magnitude of stimulus-driven behavioral modulation and the optimal (best) FM-stimulus phase were consistent across sessions within an individual.

For each session separately, hit rates were calculated for each FM-phase bin (**Fig. 4a**). Then, we fitted a cosine function to hit rates as a function of phase for each participant and each session. The resulting amplitude parameter from the cosine fit was taken as an index of the strength of the behavioral modulation. Compared to surrogate datasets, significant behavioral modulation was observed for 31/38 participants in session 1 and 34/38 in session 2 (p < 0.05, Bonferroni corrected, **Fig. 4b**).

**Figure 4.**
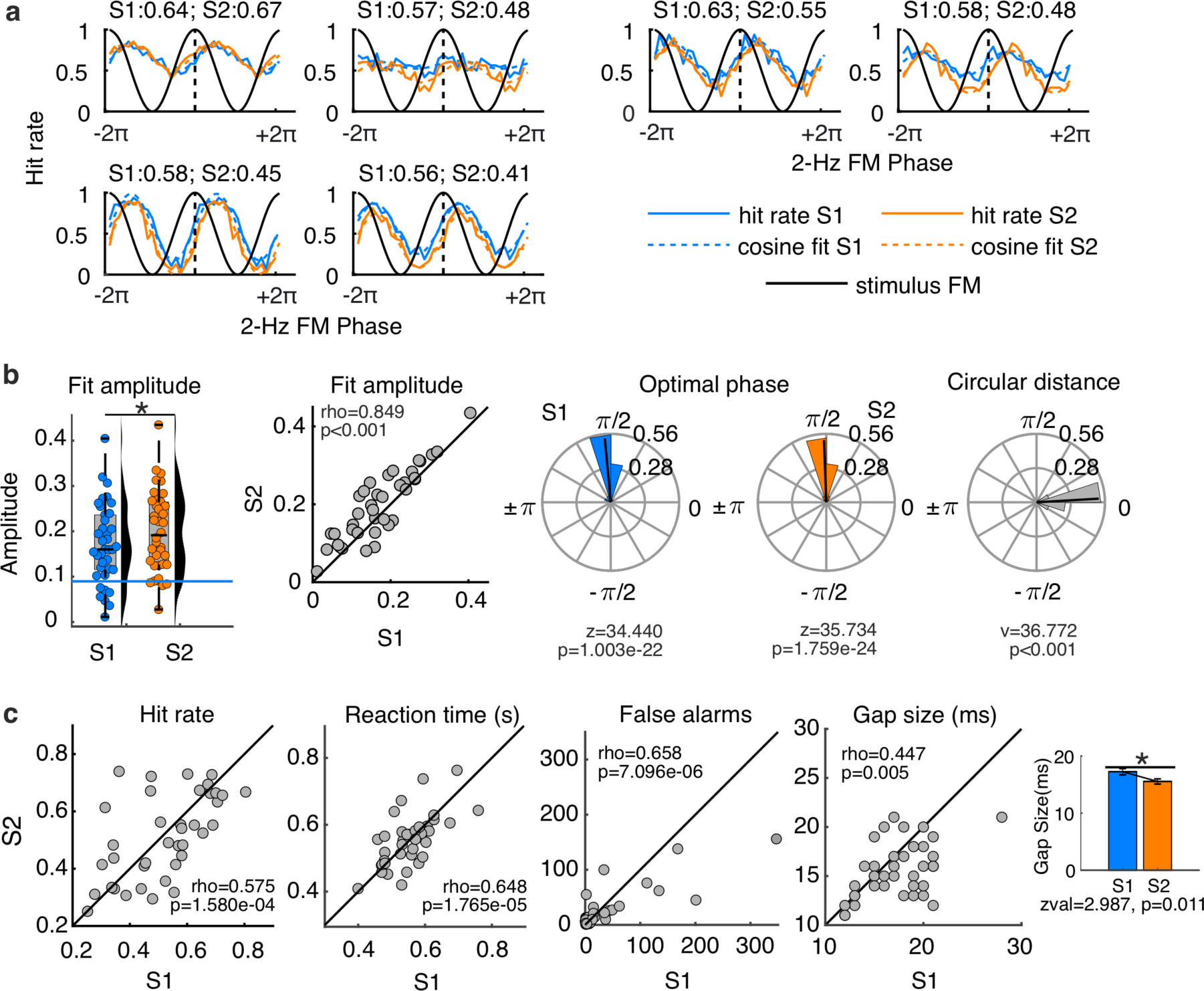
Stimulus-driven behavioral modulation and its reliability. (a) Hit rates as a function of 2-Hz FM stimulus phase. Dashed blue and orange lines represent the fitted cosine functions for sessions 1 and 2, respectively. Numbers on top of each graph show the mean hit rate across phase bins for each session. Each graph shows data for a different participant. In all panels, blue represents session 1 (S1) and orange represents session 2 (S2). (b) The plot on the left shows the distribution and spread of the amplitude of the observed sinusoidal modulation for each session. The box plot shows median (black horizontal line), 25th and 75th percentiles (box edges) and extreme datapoints (whiskers). Each circle represents a single participant. Horizontal blue and orange lines represent the 0.999 quantile (corresponding to p < 0.05, *Bonferroni* corrected) from the distribution of amplitude parameters from the surrogate datasets, for session 1 and session 2 respectively. Scatter plot in the middle shows the correlation between the fit amplitudes for the 2 sessions. Solid line is the diagonal. Circular histograms on the right of the panel show individual optimal phases (i.e., FM-stimulus phase with highest hit rate in the cosine fit) separated by session (left and middle histograms) and the circular distance between the two (right histogram). Z and P-values in the plots refer to the results from the Rayleigh test. V and p-value for the inter-session circular distance corresponds to the V test results. (c) Scatter plots show inter-session correlations for hit rates, reaction times, false alarms rates, and gap durations. Solid lines are the diagonal. *p = 9.69e-04. Distribution plots were created using a modified version of the toolbox *distributionPlot* by Dorn J, 2008.

The fitted modulation amplitude values were higher in session 2 compared to session 1 (z = 3.299, p = 9.694e-04), however, they were also highly correlated between sessions (rho = 0.849, p <0.001, **Fig. 4b**), indicating that FM-induced behavioral modulation was reliable, but increased across sessions overall. Individual optimal FM phases (the FM-stimulus phase yielding highest performance) were estimated from the fitted cosine functions per participant and per session (**Fig. 4b**). Optimal phases were not uniformly distributed, but clustered in one half of the FM cycle (Rayleigh test; session 1: z = 34.440, p = 1.003e-22; session 2: z = 35.734, p = 1.759e-24). Optimal phases were also highly correlated between sessions (circular–circular correlation; rho = 0.620, p = 0.036, **Fig. 4b**). Moreover, circular distance between optimal phases in session 1 and session 2 was clustered around zero (V test; v = 36.772, p <0.001), suggesting that optimal phase is also a reliable attribute of FM-induced behavioral modulation.

We recognized that high reliability in optimal FM phase across sessions may have been at least partially attributable to clustering of optimal phases across participants in the first place – since optimal phases were significantly clustered across participants, we wanted to be careful not to overinterpret similar optimal phase across sessions as reflecting within-participant reliability. In order to test this possibility, we conducted a permutation test in which the order of participants in session 2 was permuted 1000 times relative to the session-1 order, and the circular–circular correlation between individual optimal phases in session 1 and (permuted) 2 was computed. If inter-session correlations are driven by the similarity between participants, permuting the participant order in session 2 should not have affected inter-session correlations. However, demonstrating the reliability of optimal phase, the inter-session correlation in the original data was higher than all values in the distribution of inter-session correlations from the permuted samples (p <0.001).

In addition to the fitted amplitude parameters and optimal FM phases for gap-detection performance, hit rates (over the entire session), false-alarm rates, reaction times, and threshold gap durations were also highly correlated across sessions (all rho ≥ 0.447, p ≤ 0.005, **Fig. 4c**). When each of the dependent measures was directly compared across sessions, a significant difference was observed only for the gap duration (*Wilcoxon signed rank test*, z = 2.987, p = 0.011, *Bonferroni* corrected). Individually adjusted gap durations estimated using our threshold procedure were shorter in session 2 than in session 1. Since hit rates were not significantly different between sessions, the decrease in threshold gap duration suggests that participants experienced some degree of learning or practice effect, and could recognize shorter gaps in session 2. The reduction in threshold gap duration between session 1 and 2 was significantly correlated with music perceptual abilities (rho = 0.385, p = 0.017), as measured with the Goldsmiths Musical Sophistication Index (Gold-MSI, (Mullensiefen et al., 2014)), very tentatively suggesting that individuals with stronger music skills might have experienced a greater benefit of repeated exposure across sessions.

### Pre-stimulus neural 2-Hz phase and alpha amplitude predict gap detection

In the previous sections, we showed that the 2-Hz FM stimulus entrained neural activity at the modulation frequency and that gap-detection performance was sinusoidally modulated by FM phase. Therefore, we expected that pre-gap brain activity should also predict gap-detection performance. We examined the effects of pre-gap neural phase and amplitude in the FM stimulus frequency band (2 Hz), pre-gap neural phase and amplitude in the alpha frequency band (7-12 Hz), and their interactions, on gap detection performance (see Methods). Both entrained 2-Hz and non-entrained alpha activity were taken into account since our initial FFT analysis showed stimulus-driven modulation of both, and because it has previously been shown that entrainment lapses are accompanied by increases in alpha amplitude (Lakatos et al., 2016), which led us to the hypothesis that both delta and alpha frequencies might contribute to gap detection (**Fig. 2a-b**). Prior to averaging 2-Hz phase and alpha phase across the relevant electrodes within each cluster, single trial phase values were realigned to each electrode’s reference phase (see Methods). This was implemented because averaging over electrodes can lead to phase cancellation and imprecise estimation of phase effects on performance. Fourteen mixed-effects logistic regression models were fitted to the data using different combinations of regressors aiming to predict trial-based gap detection performance (hit/miss). In both sessions, the winning model (smallest AIC values) included the predictors pre-gap neural 2-Hz phase and pre-gap alpha amplitude (**Fig. 5a**; winning model vs. next best model, session 1: ΔAIC = 1.682, LRStat = 14.318, ΔDF = 8, p = 0.074; session 2: ΔAIC = 2.870, LRTstat = 13.13, ΔDF = 8, p = 0.107). While ΔAIC between the winning model (2-Hz phase + alpha amplitude) and the next best model (2-Hz phase + alpha amplitude + alpha phase) was smaller than 2 in session 1, we decided to take this model because 1) it showed the lowest AIC in both sessions, 2) it was the simplest model of the two and 3) the LRT suggested that the more complex model was not better at fitting our data. Adding the predictors 2-Hz amplitude, alpha phase, or the interactions 2-Hz phase*alpha amplitude and 2-Hz phase*alpha phase produced higher AIC values. **Table 1** shows the AIC and ΔAIC values for all tested models.

**Figure 5.**
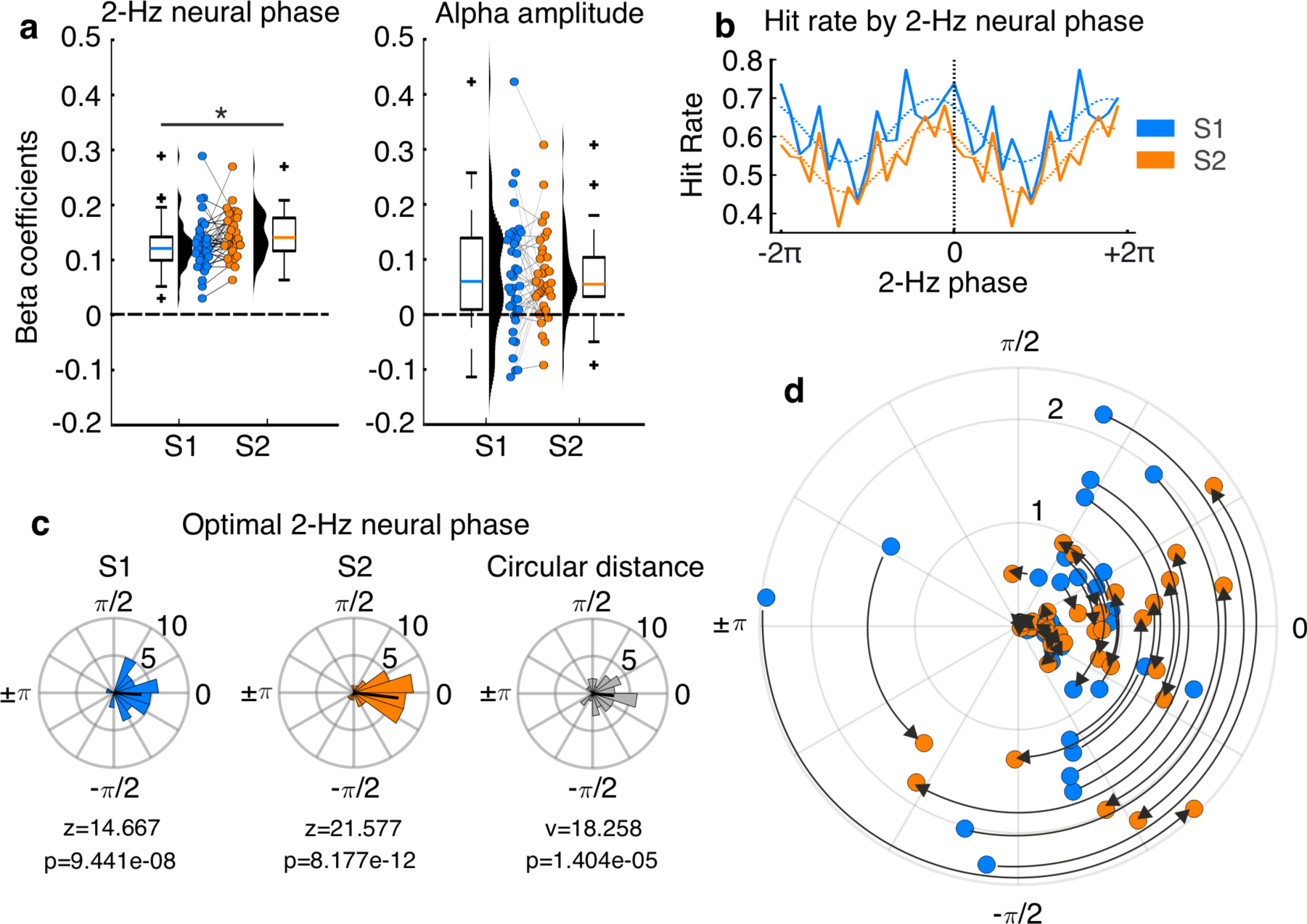
Effect of pre-gap neural activity on gap detection. (a) Beta estimates (including distribution and spread) for 2-Hz phase (left) and alpha amplitude (right) from the winning mixed-effects logistic regression models fitted to the EEG data. Box plots show median (horizontal line), 25th and 75th percentiles (box edges) and extreme datapoints (whiskers). Black crosses represent outlier values. Each circle represents a single participant. Horizontal dashed lines mark the zero line. (b) Hit rates as a function of pre-gap 2-Hz neural phase for one representative participant. Solid lines represent actual data and dashed lines represent the cosine fits. Two cycles are concatenated for visualization purposes. (c) Circular histograms show the distribution of optimal neural 2-Hz phases across participants for each session (left/middle graphs) and the circular distance between both sessions (right). Z and p-values in the plots refer to the results from the Rayleigh test. V and p-value for the inter-session circular distance corresponds to the V test results. (d) Optimal 2-Hz neural phase for session 1 (blue) and session 2 (orange). Black arrows show the phase shift from session 1 to session 2. The Radius in the plot represents the absolute circular distance between sessions. Each circle represents a single participant and session. S1: session 1 (blue); S2: session 2 (orange). *p = 0.03, *Bonferroni* corrected.

To test whether the winning model had a classification accuracy higher than chance, the ROC and corresponding AUC were calculated. AUC values were then compared to the distribution of AUCs obtained when fitting the same model to surrogate data (see Methods). For both sessions, the winning model showed higher AUC than all values in the surrogate distribution, demonstrating that pre-gap neural 2-Hz phase and alpha amplitude predicted gap detection accuracy significantly better than chance (AUC S1 = 0.675, S2 = 0.677, p <0.001).

Beta coefficients for 2-Hz phase effect in session 2 were higher than in session 1 (z = 2.444, p = 0.029, *Bonferroni* corrected, **Fig. 5a**). No significant difference between sessions was observed for the effect of alpha amplitude (z = 0.051, p = 0.960, uncorrected). For most participants, beta coefficient for the effect of alpha amplitude were positive in both sessions, indicating that alpha amplitude was higher for hits than for misses.

Next, we evaluated the clustering across participants of optimal neural 2-Hz phase, both within and across sessions. Trials were sorted into 18 equally spaced bins according to the realigned instantaneous neural 2-Hz phase (**Fig. 5b**). Hit rates were calculated for each bin and cosine functions were fitted to each individual participant’s data in order to estimate optimal neural phase, similar to the behavioral analysis. In both sessions, optimal 2-Hz neural phases were significantly clustered across participants (all z ≥ 14.667, p ≤ 9.441e-08, **Fig. 5c**). Circular distances between optimal phases for both sessions were also clustered around zero (v = 18.258, p = 1.404e-05, **Fig. 5c**). Unexpectedly, the circular– circular correlation for optimal phases between sessions was not significant (rho = -0.124, p = 0.429). Plotting session-wise optimal phase shifts suggested that while mean phase distances were around zero, optimal phase for different participants changed but in different directions (**Fig. 5d**). Interestingly, the absolute optimal-phase distance between sessions correlated with the overall (mean) magnitude of the 2-Hz phase effect on behavior (beta coefficients; rho = -0.339, p = 0.038): participants with strongest phase effect showed higher optimal phase consistency across sessions. No significant association between phase distance and inter-session interval or time of the day was observed (all |rho| ≤ 0.216, p ≥ 0.193, uncorrected).

### Pre-gap 2-Hz phase predicts alpha amplitude

In the previous section, we showed that both 2-Hz phase and alpha amplitude predict single-trial gap detection performance. However, it is not clear whether they represent two independent mechanisms or if the two neural signatures might be coupled. A natural question here would then be whether there is a significant phase–amplitude coupling between 2-Hz and alpha activity. To answer this question, a generalized mixed effects model was implemented with single-trial pre-gap alpha amplitude as the dependent variable and pre-gap 2-Hz phase as the predictor (sine and cosine), using both fixed effects terms as well as random intercept and slope. We expected that if 2-Hz phase and alpha amplitude are coupled, 2-Hz phase should significantly predict alpha amplitude. To test for the significance of the phase–amplitude relationship relative to chance, beta estimates for each session were compared to the beta estimates obtained from fitting the same model to surrogate datasets, which were created by shuffling single-trial alpha amplitudes while keeping 2-Hz phase fixed. For both sessions, beta estimates from the real model were higher than all values in the distribution of beta estimates from the surrogate data (p <0.001, **Fig. 6**), indicating that pre-gap 2-Hz neural phase significantly predicted pre-gap alpha amplitude.

**Figure 6.**
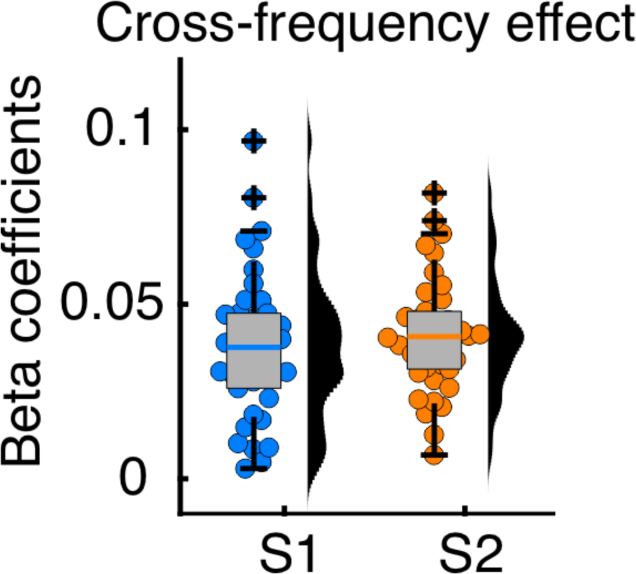
Predicting alpha amplitude from 2-Hz neural phase. The graph shows the distribution and spread of the beta coefficients from the generalized linear mixed effects model predicting pre-gap alpha amplitude from pre-gap 2-Hz neural phase (left). Box plots show median (colored horizontal lines), 25th and 75th percentiles (box edges) and extreme datapoints (whiskers). Each circle represents a single participant. S1 (blue colors): session 1, S2 (orange colors): session 2.

### Control Experiment: Gap detection is modulated by the FM rate

Here, we found that optimal FM phases were clustered in one half of the FM cycle (although we did observe some variability across participants). This was somewhat surprising because previous studies using the same task and very similar stimuli reported that optimal phases were uniformly distributed around the FM cycle (Henry and Obleser, 2012). One major difference between the two studies was the FM frequency (rate): 3 Hz in the previous study (Henry and Obleser, 2012) and 2 Hz in the current study. We hypothesized that individual variability in stimulus-driven behavioral modulation might be influenced by the FM frequency: slower FM frequencies (in this case 2 Hz) should be easier to track and therefore participants should synchronize to such stimuli more similarly, while individual differences are more pronounced for higher FM frequencies, which might be more difficult to track and therefore result in phase slips.

We tested this hypothesis in a control experiment, where 25 participants performed the same gap-detection task as in the main experiment, but FM rate varied between blocks and took on values of 1.5 Hz, 2 Hz, 2.5 Hz, or 3 Hz (see Methods, **Fig. 7**). Overall hit rates were significantly modulated by FM rate (repeated measures ANOVA, F (3,72) = 13.374, p = 4.980e-07); lower hit rates were observed for higher FM rates (**Fig. 7a**). Note that here, overall hit rates are a useful performance measure, because we did not titrate gap duration to match hit rates across FM rates or across individuals. FM rate also significantly affected the amplitude of the stimulus-induced behavioral modulation, albeit in opposite direction to the hit-rate effect; amplitude values increased as a function of FM rate (F (3,72) = 6.336, p = 7.128e-04, **Fig. 7b**)). To test whether the clustering of individual optimal phases was dependent on the FM rate, individual optimal phases were estimated from cosine fits (**Fig. 7c**), and the resultant vector length across participants was calculated for each FM rate (**Fig. 7c**). For this calculation, individual optimal phases were weighted using the amplitude parameter of the fit. We did this because estimation of optimal phase is less accurate the lower the amplitude parameter (i.e., weaker sinusoidal modulation of behavior), therefore we reduced the weight of optimal phases from low-amplitude fits in the resultant vector calculation. A linear model was used to predict resultant vector length from FM rate and an intercept parameter. A significant fit for the linear term would suggest that indeed, phase clustering significantly decreased (or increased) with increasing FM rate. Significance was evaluated using permutation tests. The permutation distribution was created by 1) computing resultant vector lengths over 1000 iterations (in each iteration the FM rate labels where permuted for each participant) and fitting the same linear model to the simulated vector lengths as for the original data. The *t-statistic* for the effect of FM rate on vector length in the original data was compared to the distribution of *t-statistics* obtained from the permuted datasets. Contrary to our prediction, the phase clustering increased with increasing FM rate (*t-stat* = 4.419, permutation p = 0.047). In sum, FM rate affected hit rates, amplitude of the stimulus-induced sinusoidal modulation, as well as optimal phase clustering across participants.

**Figure 7.**
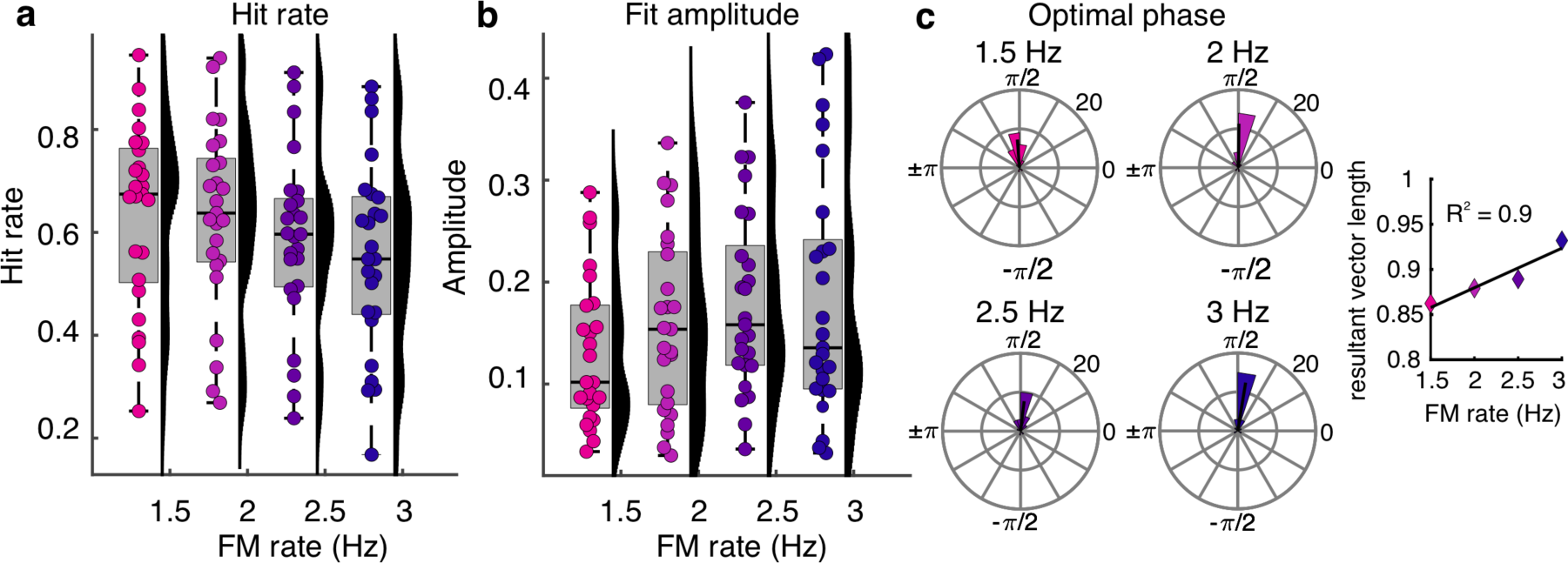
Control experiment. (a) The graph shows the distribution and spread of individual hit rates as a function of stimulus FM rate. Box plots show median (black horizontal line), 25th and 75th percentiles (box edges) and extreme datapoints (whiskers). Each circle represents a single participant. (b) Same as in (a) but for the amplitude of the sinusoidal modulation. (c) Circular histograms showing the distribution of optimal FM-stimulus phases for each FM rate. The inset on the right shows the resultant vector length computed across individual optimal phases separated by FM rate.

### Gap detection is also influenced by stimulus carrier and time of occurrence

Although we tried to minimize the possibility that the FM phase effects may have unknowingly been driven by acoustic confounds, we nonetheless tested which stimulus characteristics beyond FM phase influenced gap-detection hit rates. Fitting different mixed-effects logistic regression models, we examined the extent to which single-trial gap detection accuracy in the main experiment was influenced by 1) FM phase, 2) global experiment time (when the gap occurred over the whole experiment, quantified as the gap number within a session), 3) local time (when the gap occurred within a stimulus), 4) the center carrier frequency of the stimulus in which the gap was presented, 5) the center frequency at gap onset, 6) cochlear-scaled entropy at gap onset, 7) the interaction of FM phase with global time, quantifying the extent to which stimulus-driven behavioral modulation changed over the course of the experiment, and 8) with local time, quantifying the extent to which stimulus-driven behavioral modulation changed over the course of each individual stimulus. Each predictor was modeled as a fixed factor with a random intercept and slope. The winning model was chosen based on AIC and the LRT. To assess the winning model’s significance relative to chance, surrogate datasets were created by permuting, for each participant, accuracies (1/0) across trials while keeping all other parameters the same. This procedure was repeated for 1000 iterations. The same model was fitted to each permuted dataset. ROC and AUC were computed for the real model and those fitted to the permuted datasets. The model was considered to be significant if the AUC values was above 95% percent of the AUC values from the permuted samples, for a corresponding p value of 0.05.

For session 1, the winning model included FM stimulus phase, global time, local time, main stimulus carrier and the interactions of phase * global time and phase * local time (**Fig. 8**, **Table 2**). In contrast, for session two, the best model did not include any interaction term but only main effects FM stimulus phase, global time, local time, and main stimulus carrier (**Fig. 8**).

**Figure 8.**
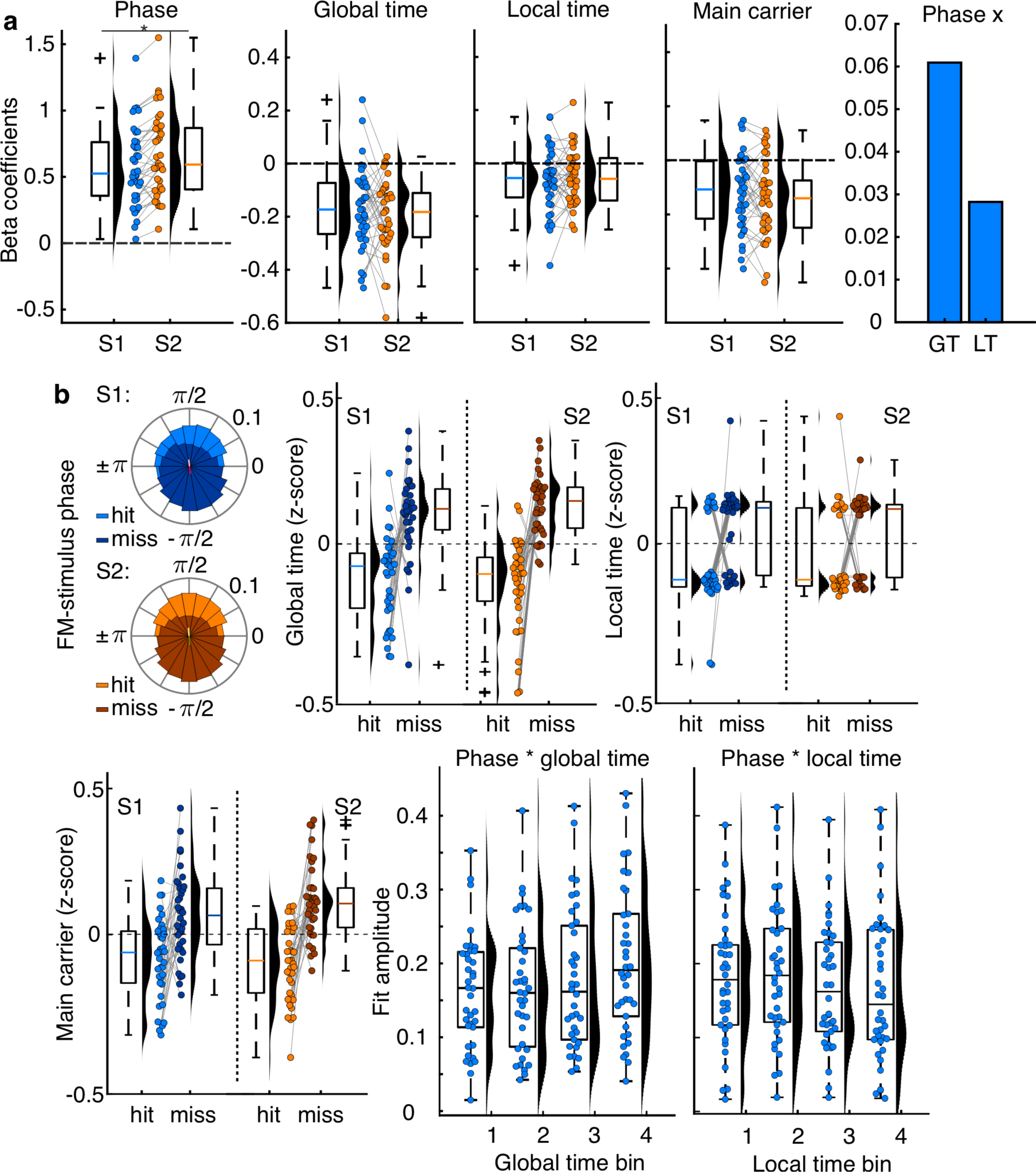
Effect of stimulus properties on gap detection. (a) Distribution plots showing the beta estimates resulting from mixed effect logistic regression models. Box plots show median (horizontal solid black line), 25^th^ and 75^th^ percentiles (box border), extreme values (whiskers) and outliers (black cross). (b) Visualization of the main effects shown in (a), i.e., difference of hits vs misses in terms of FM-stimulus phase, global time, local time and stimulus center carrier frequency. The two plots in the bottom right show the interaction phase * global time and phase * local time. In the plots, the amplitude of the sinusoidal modulation (fit amplitude) is shown after fitting cosine function to hit rates by FM-stimulus phase independently to four global/local time bins. S1: session 1; S2: session 2. * p = 0.006, *Bonferroni* corrected.

Mean beta values for global time, local time, and main carrier frequency were negative, indicating that gap detection performance decreased with increasing global time, local time, and main carrier frequency (**Fig. 8a**). That is, gap detection was better earlier in the session as would be expected from fatiguing over the experiment, earlier in each stimulus, and for relatively low carrier frequencies (**Fig. 8b**). Interestingly, when we plotted the local-time effect, we realized that it arose from two distinct groups of participants: a bigger group who better detected gaps early in the stimulus, and a smaller group who better detected gaps later in the stimulus (**Fig. 8b**). To visualize the interaction effects observed in session 1, trials were grouped in four bins according to their time of occurrence within the session (global time) or within each 20-s stimulus (local time). Hit rates were computed for each FM phase and time bin, and cosine functions were fitted to each bin’s data. Visual inspection of this analysis showed that the amplitude of the stimulus-driven behavioral modulation was higher later in the session and earlier within the stimulus (**Fig. 8b**). Beta estimates for the effect of FM stimulus phase were significantly higher in session 2 compared to session 1 (z = 3.836, p = 5.005e-04, *Bonferroni* corrected). No significant difference between sessions were observed for any of the other predictors (|z| ≤ 1.240, p ≥ 0.215, uncorrected).

As already mentioned, additional mixed effects logistic regression models were fitted to the individual data using other combination of parameters, including cochlear-scaled entropy and/or the carrier frequency at the exact time the gap was presented, taking into account the sinusoidal modulation. All such models showed higher AIC values than the winning model (**Table 2**, winning model vs. next best model, session 1: ΔAIC = 2.520, LRTstat = 5.480, ΔDF = 4, p = 0.241; session 2: ΔAIC = 4.025, LRTstat = 3.975, ΔDF = 4, p = 0.409). Interestingly, the model with the highest AIC (and therefore worst performance; ΔAIC relative to winning model; session 1: > 2483.513; session 2: > 3200.484, see **Table 2**) was the one including acoustic parameters (including gap onset carrier frequency) but excluding FM stimulus phase information, suggesting that in fact, FM phase was the best predictor of gap detection accuracy. For both sessions, the winning model had AUC values higher than all values in the distribution of surrogate datasets, proving a classification accuracy higher than chance (AUC; S1 = 0.744, S2 = 0.758; p <0.001).

## Discussion

In the present study, we investigated the test–retest reliability of neural entrainment and its relevance for auditory perception. Participants detected silent gaps embedded in 2-Hz FM stimuli while EEG activity was recorded in two separate sessions. We showed that: 1) neural activity was entrained by the FM stimuli, and perception was modulated in a sinusoidal manner by both stimulus and brain phase; 2) both entrainment strength and the magnitude of stimulus-driven behavioral modulation were reliable across sessions; 3) pre-gap delta phase and alpha amplitude predicted moment-by-moment fluctuations in performance.

### Neural entrainment is reliable over time

Over the past years, a great deal of attention has been paid to entrainment, and arguments range from the existence of entrainment per se (Rimmele et al., 2018; Obleser and Kayser, 2019; Meyer et al., 2020) to its role for auditory perception specifically (Ng et al., 2012; Sameiro-Barbosa and Geiser, 2016; Lakatos et al., 2019). Neural entrainment to rhythmic stimuli has been observed across a range of sounds, including simple stimuli such as tone sequences, amplitude-modulated or frequency-modulated sounds, and more complex auditory stimuli such as music and speech (Henry and Obleser, 2012; Doelling et al., 2014; Henry et al., 2014; Doelling and Poeppel, 2015; Lakatos et al., 2016; Henry et al., 2017; Ten Oever et al., 2017; Riecke et al., 2018). However, until now, little attention has been paid to the reliability of neural entrainment. This is despite the fact that the reliability of entrainment is of paramount importance both for understanding its contribution to auditory perception and for developing effective targeted interventions. Here, we showed that neural entrainment to FM sounds is reliable over time: we observed high inter-session correlations for metrics of entrainment strength, such as the spectral amplitude of neural activity at the stimulus frequency, and perhaps more importantly, the phase lag between stimulus and brain. Interestingly, while no differences in either stimulus–brain phase lag or optimal stimulus phase were observed between sessions, both the effect of FM-stimulus phase and the effect of pre-gap 2-Hz neural phase on gap detection were stronger in session 2 than in session 1. This suggests that while highly reliable, the degree to which behavior is influenced by auditory rhythms can potentially be trained.

### Entrained delta and ongoing alpha activity influenced gap-detection performance

Here, we observed that trial-by-trial gap-detection performance was mainly predicted by pre-gap neural 2-Hz phase and alpha amplitude. These results are in line with previous studies showing that the phase of neural oscillations prior to target occurrence predicts perception in different sensory domains (Hanslmayr et al., 2007; Vanrullen et al., 2011; Mathewson et al., 2012; Weisz et al., 2014; Wostmann et al., 2019a). For example, enhanced detection or faster reaction times have been reported for sensory stimuli presented at the optimal phase of delta (Stefanics et al., 2010), theta (Busch et al., 2009; Busch and VanRullen, 2010), or alpha oscillations (Busch et al., 2009; Vanrullen et al., 2011). Our results also showed that, in a rhythmic listening context, gap-detection performance was not just a product of the entrained delta activity but also depended on non-entrained (ongoing) alpha activity. Comodulation of behavior by different frequencies has previously been reported (Henry et al., 2014). Critically, such comodulation was specific to the entrained frequencies, which led the authors to conclude that environmental rhythms reduce dimensionality of neural dynamics. Here we expand this view by showing that both entrained and ongoing brain oscillations could potentially comodulate behavioral performance. We interpret the contributions of both entrained and ongoing activity to perception as potentially reflecting an interplay between stimulus driven (sensory, bottom-up) and internally driven (top-down) processes.

Although our paradigm was not designed to provide a time-resolved look at the interplay between delta and alpha oscillations, we observed that single-trial 2-Hz pre-gap phase predicted single-trial alpha amplitudes, which we interpret as delta–alpha phase-amplitude cross-frequency coupling. In the current context, we are unable to tease apart the possibilities that, on the one hand, entrained delta oscillations drove alpha-amplitude fluctuations, or on the other hand, the rhythmic auditory stimulus drove both delta oscillations as well as alpha-amplitude fluctuations, but that the two neural signatures are functionally independent. At first glance, the observed delta-alpha coupling seems to contradict the finding that both delta-phase and alpha amplitude predict gap detection: if delta-alpha coupling is perfect, including only one of the two predictors should be enough to explain gap detection. The fact that the winning model includes both predictors however indicates that delta-alpha coupling is not perfect. Our time-resolved ITPC analysis showed that coherence (entrainment) at the stimulus frequency fluctuates over time (Lakatos et al., 2016). We speculate that delta-alpha coupling is higher when delta activity successfully entrains to the FM stimulus compared to when entrainment is poor. The contribution of alpha amplitude to gap detection might then be stronger in those trials. Therefore, both predictors are necessary in the winning model. In other words, our results are consistent with alternating influences that might occur as a result of “lapses” of entrainment during which alpha oscillations might have a stronger effect on perception (Lakatos et al., 2016). It is possible that when entrainment is high, auditory perception is mostly modulated by the entrained activity, however, during entrainment lapses, perception is shaped by internal ongoing activity. Or conversely, participants could have adopted the strategy of trying to ignore the rhythm in order to perform the task, but as their alpha–attention system lapsed, they were forced into a rhythmic–entrainment mode.

Quite a bit of previous work has demonstrated the importance of alpha activity for “gating” near-threshold stimuli into awareness (Mathewson et al., 2009; Mathewson et al., 2012), which goes beyond the idea of alpha activity indexing lapses of entrainment. However, the vast majority of these studies have been conducted in the visual modality (Spaak et al., 2014; Fodor et al., 2020; Hutchinson et al., 2020), and it has been argued that alpha activity does not contribute to near-threshold auditory perception (VanRullen et al., 2014). Thus, one open question relates to the precise role of alpha oscillations in shaping near-threshold auditory perception, in particular in a rhythmic auditory context. Answering this question requires respecting the observation of different alpha rhythms with different neural generators (Isaichev et al., 2001; Weisz et al., 2011; Billig et al., 2019; Keitel et al., 2019). Moreover, several studies linking alpha oscillations to attention have suggested that alpha oscillations could play at least two different roles: i.e., a facilitatory role where it can enhance target processing, or a suppressive role where alpha activity can suppress the processing of distractors (Bonnefond and Jensen, 2012; Wostmann et al., 2019b; Schneider et al., 2020). We hypothesize that the alpha effect observed in our study is reflecting something like distractor suppression. While delta oscillatory activity entrained to the FM stimulus facilitates target processing at the optimal delta phase, we speculate that ongoing alpha activity might play a role in suppressing the distracting stimulus itself (complex noise), in an attempt to also maximize target detection. In this case, gap detection performance would be predicted by both mechanisms, in line with the current results.

### Behavioral entrainment depends on modulation rate

In the current study, we were surprised to observe that the optimal *stimulus* phase that yielded best gap-detection performance was consistent across participants. In previous work, optimal stimulus phase was uniformly distributed across participants, and this observation was critical to our argument that stimulus-driven behavioral modulation was the result of neural entrainment and not an artifact of stimulus acoustics (Henry and Obleser, 2012). One main difference between the current study and previous work was the FM rate, which was slower here (2 Hz) than in previous work (3 Hz). This led us to conduct a control experiment where we examined gap-detection performance for stimuli with FM rates varying between 1.5 and 3 Hz (in 0.5-Hz steps). We observed that hit rates and the amplitude of the stimulus-driven behavioral modulation were significantly modulated by FM rate. Contrary to our initial hypothesis, we found that, when controlling for the modulation amplitude, optimal phase became more consistent as FM rate increased. Such findings were unexpected to us and suggest that while FM rate plays a major role on behavioral entrainment to FM stimuli, it does not explain the difference in phase clustering between the current study and (Henry and Obleser, 2012). In addition to, and perhaps more importantly than, the FM rate, the two studies also used different FM-modulation depths: 67% (center-to-peak) here and 37.5% in (Henry and Obleser, 2012). It remains to be tested in future studies whether such modulation depth influences phase clustering across participants or not.

## Conclusion

Taken together, our results showed that FM stimuli entrained neural activity and sinusoidally modulated near-threshold target detection: both signatures of entrainment as well as its behavioral consequences were reliable across sessions. This demonstration is a critical prerequisite for research lines focused on targeted interventions for entrainment but has to our knowledge gone untested until now. Moreover, gap-detection performance was predicted by entrained neural delta phase and ongoing alpha amplitude, suggesting that delta phase and alpha amplitude underpin different but potentially simultaneously active neural mechanisms and together shape perception.

## Acknowledgments

This work was supported by a European Research Council Starting Grant (BRAINSYNC) and a Max Planck Society Research Group granted to MJH. We thank Nicole Huizinga for helping with the data collection. We thank Hanna Kadel, Dominik Thiele, and Johannes Messerschmidt for technical support and for helping with the EEG preparation and participants recruitment. We also thank Cornelius Abel for technical support.

## Notes

### Competing Interest Statement

The authors have declared no competing interest.

